# Brain-derived neurotrophic factor and adenosine A2A receptor interaction modulates oligodendrogenesis derived from postnatal SVZ neural stem cells

**DOI:** 10.1101/2024.09.26.615137

**Authors:** J.M. Mateus, A. Barateiro, B. Santos, J.B. Moreira, N. Plachtij, A.M. Sebastião, A. Fernandes, S. Xapelli

## Abstract

Oligodendrocytes (OLs) are vital for myelin formation in the Central Nervous System. OLs can be produced by the maturation of oligodendrocyte precursor cells (OPCs) present throughout the brain parenchyma or from the differentiation of subventricular zone-derived neural stem cells (SVZ-NSCs). Importantly, efficient differentiation from SVZ-NSCs remains a significant area of research due to its potential for remyelination in demyelinating disorders. In this work, we studied the role of brain-derived neurotrophic factor (BDNF) and adenosine A2A receptors (A2ARs), as well as the putative crosstalk between these two modulatory mechanisms, in regulating oligodendrogenesis from SVZ-NSCs. Using a neurosphere culture system, we observed that BDNF significantly increased the mRNA expression levels of OPC cell markers after 2 days in vitro (DIV), an effect blocked by the A2AR antagonist ZM 241385. This early transcriptional regulation by BDNF was followed by changes in the percentage of both OPCs and mature OLs in culture at DIV 7 and 14. Interestingly, blocking A2ARs prevented the potentiating effect of BDNF on the percentage of OLs at DIV 14. Concerning the morphology of mature OLs, BDNF influenced their maturation by reducing branching near the soma at DIV 7, an effect that was not observed at 14 DIV, when all treatments resulted in similar OL morphology. Overall, our results establish BDNF as a regulator of OL formation from SVZ-NSCs, with A2AR-BDNF interaction modulating the differentiation process.

## Introduction

In the adult mammalian brain, neural stem cells (NSCs) are located in two regions, known as neurogenic niches: the subventricular zone (SVZ), a thin layer of cells lining the lateral ventricles, and the subgranular zone (SGZ) of the hippocampal dentate gyrus (DG)[1]. NSCs maintain their abilities of multipotency and self-renewal. However, in vivo studies have shown that NSCs from the SGZ can give rise to only neurons and astrocytes[2], whereas SVZ-derived NSCs (SVZ-NSCs) can differentiate into the three types of neural cells: neurons, astrocytes and oligodendrocytes[3–5].

Oligodendrocytes (OLs) are the cells responsible for myelin formation in the Central Nervous System, playing a crucial role under demyelinating conditions. Importantly, studies have shown that both parenchymal oligodendrocyte precursor cells (OPCs) and those derived from SVZ-NSCs can differentiate into OLs and partially repopulate lesioned regions[6–8]. Nevertheless, the remyelination process often remains incomplete, highlighting the need to find potential modulators that promote the formation of functional mature OLs capable of repairing the damaged myelin and restoring neuronal function as disease progresses.

Brain-derived neurotrophic factor (BDNF), a neurotrophin highly expressed in the mammalian brain, has been widely studied for its role in modulating OPC survival and myelination[9, 10]. Notably, work using adult mouse SVZ-derived neurospheres has shown that low BDNF concentrations promote self-renewal and proliferation of NSCs via the TrkB receptor, while higher BDNF concentrations, which also bind the p75NTR receptor, enhance TrkB-dependent self-renewal and proliferation and also promote NSC differentiation[11]. In BDNF heterozygous mice, the role of BDNF in oligodendroglial cells is both transient and limited to specific regions during postnatal development[12]. Furthermore, BDNF-TrkB signalling plays a major role in myelination during development, as could be concluded in a study using a conditional knockout model of TrkB receptors [13]. Interestingly, this study also showed that OPC proliferation is potentiated in conditional TrkB knockout mice[13], while another study showed that BDNF potentiates OPC proliferation in cuprizone-induced demyelinating conditions[14]. These data suggest that BDNF actions upon OPC proliferation may be different in non-pathological and in pathological demyelination conditions. In line with the evidence for a relevant role of BDNF in demyelinating diseases is the report that brain delivery of BDNF in mice with experimental autoimmune encephalomyelitis, the most commonly used animal model of MS, promotes neuroregeneration by activating OLs as well as enhancing remyelination[15].

Several studies have highlighted the role of adenosine and adenosine A2A receptors (A2AR) in gliogenesis and neurogenesis under healthy and pathological conditions[16]. Our group has previously shown that activation of A2ARs in postnatal rat DG-derived neurospheres promotes BDNF release, and that extracellular BDNF is required for A2AR actions[17]. Interestingly, A2AR activation has been shown to promote BDNF effects on hippocampal synaptic transmission, with A2AR antagonists blocking this effect[18]. The interaction between these two regulatory mechanisms has been studied mostly focusing on the hippocampus and/or upon synaptic signalling, but its role in SVZ-derived oligodendrogenesis remains to be studied.

Clarifying how BDNF modulates OL formation from SVZ-derived NSCs and understanding the impact of A2AR blockage on this process, can offer valuable insights into new potential targets for in vivo modulation. In this study, using SVZ-derived neurospheres we observed that BDNF promoted early OPC marker expression and later enhanced OPC and OL differentiation, along with inducing stage-specific morphological changes in OLs. Interestingly, A2AR inhibition modulated BDNF effects in mature OLs. Taken together, our results start to unveil how the interactions between BDNF and A2ARs can influence oligodendroglial differentiation from postnatal SVZ-NSCs. This knowledge may be used to promote OL formation and enhance remyelination in demyelinating conditions.

## Materials & Methods

### 1. Ethics

All procedures were conducted in accordance with the European Directive 2010/63/EU and the Portuguese law Decreto-Lei 113/2013 for the protection of animals used for scientific purposes. The protocol was approved by the “iMM’s Institutional Animal Welfare Body” – ORBEA-iMM and the National competent authority – Direção Geral de Alimentação e Veterinária (DGAV). The pups were handled according to standard and humanitarian procedures to reduce animal suffering.

### 2. Animals

C57BL/6J males were kept in individual standard housing and females were grouped in pairs. All animals were kept on a 12 h light/dark cycle with food and water *ad libitum*. Breeding attempts were only started after animals reached sexual maturity (8 weeks of age for males and 6 weeks of age for females). For maximal breeding yield, breeding trios were used until animals reached one year of age or ten successful mating sessions. All efforts were made to ensure minimal animal suffering and stress, and to use the minimum number of animals, according to standard and ethical procedures. All animals were provided with standard housing enrichment (access to disposable igloos and cardboard tubes), as well as nesting material and wooden pellets. Experiments were performed with biological material obtained from postnatal day 1 to 3 (P1-P3) C57BL/6J mice and subsequently maintained in vitro. A minimum of 3 pups was required to perform one SVZ cell culture and a minimum of 3 independent cell cultures was required for statistical analysis.

### 3. SVZ Cell Cultures

SVZ neurospheres were prepared from P1-P3 C57BL/6J mice as described in Soares *et al*., 2020[19]. Briefly, SVZ fragments were dissected out from 450 µm-thick coronal brain slices, digested with 0.05% Trypsin-EDTA (#25300054, Thermo Fisher Scientific) in Hank’s Balanced Salt Solution (#14175095, Thermo Fisher Scientific), and mechanically dissociated with a P1000 pipette. The resulting cell suspension was diluted in serum-free medium (SFM) composed of Dulbecco’s modified Eagle’s medium/Nutrient Mixture F-12 with glutaMAX (#31331028, Thermo Fisher Scientific), supplemented with 100 U/mL penicillin and 100 µg/mL streptomycin (#15070063, Thermo Fisher Scientific), 1% B27 supplement (#17504044, Thermo Fisher Scientific) and growth factors [10 ng/mL epidermal growth factor (#53003-018, Thermo Fisher Scientific) and 5 ng/mL human fibroblast growth factor (#13256-029, Thermo Fisher Scientific)] (proliferative conditions). SVZ cells were seeded in uncoated Petri dishes (60 mm diameter; #430166, Corning), and maintained for six days in a 95% air/ 5% CO_2_ humidified atmosphere at 37°C. After six days, neurospheres were plated for 24 h onto glass coverslips (12 mm diameter; #631-1577P, VWR) coated with 100 µg/mL poly-D-lysine (#P7886, Sigma-Aldrich) in SFM devoid of growth factors (differentiative conditions). After 24 h (day in vitro, DIV 0), the medium was renewed with or without (control) the pharmacological treatments (see Table 1) for the duration of the protocol. For cells kept for 14 DIV, half of the medium was replaced after 7 DIV, and pharmacological treatments were repeated.

**Table 1.** Pharmacological treatments used.

| Drug | Biological activity | Concentration used | Catalog number | Company |
| --- | --- | --- | --- | --- |
| BDNF | TrkB ligand | 30 ng/mL | NA | Regeneron Pharmaceuticals |
| ZM 241385<br>4-(2-[7-Amino-2-(2-furyl)[1,2,4]triazolo[2,3-a][1,3,5]triazin-5-ylamino]ethyl)phenol | A2AR selective antagonist | 50 nM | #1036 | Tocris Bioscience, UK |

### 4. Pharmacological Treatments

To investigate the crosstalk between BDNF and adenosine A2AR on cell viability and oligodendroglial differentiation, SVZ-neurospheres were incubated with BDNF (30 ng/mL) and a selective A2AR antagonist (ZM241385, 50 nM) (Table 1). The ligand concentrations used in the study were selected from previous published work[20]. Whenever cultures needed to be co-treated with a combination of drugs, treatment with the selective antagonist for A2AR was performed 30 min prior to the treatment with BDNF.

### 5. Semi-quantitative qReal-Time PCR

mRNA expression levels of nerve/glial-antigen 2 (NG2), platelet derived growth factor receptor α (PDGFRα), myelin basic protein (MBP) and A2AR were measured by Quantitative Real-Time PCR (qReal-Time PCR). Total RNA was isolated from SVZ-neurospheres at distinct timepoints (DIV 2, 7 and 14), using the Ribozol™ reagent method, according to the manufacturer’s instructions (VWR Life Science). After RNA extraction, the RNA concentration and purity was checked using NanoDrop ND-100 Spectrophotometer (Nanodrop Technologies). RNA was then reversibly transcribed into complementary DNA (cDNA) using Xpert cDNA Synthesis Mastermix kit (#GKB6.00100, GRiSP) according to manufacturer’s directions. Subsequently, cDNA was amplified by qReal-Time PCR on a 7300 Real-Time PCR System (Applied Biosystem) by the excitation and emission of Xpert Fast SYBR Mastermix (#GE22.2501, GRiSP). The PCR was performed in 384-well plates, with duplicates for each sample as well as a no-template control. The cycle conditions were previously optimized: 50°C for 2 min, 95°C for 10 min followed by 40 cycles at 95°C for 15 sec and 62°C for 1 min. All primers were purchased from Thermo Fisher Scientific (see Table 2 for primer sequences) and Rpl19 was used as an endogenous control to normalize the expression levels. Relative mRNA concentrations were calculated using the ΔΔCt comparative method, in which for each sample, ΔCt was determined by the difference between the Ct value of the gene of interest and the mean Ct value of Rpl19. Then, ΔΔCt value of one sample was achieved by the difference between its ΔCt value and the ΔCt of the sample chosen as reference (i.e., the control (CTRL) group for that culture). Results are presented as relative expression to the standard gene.

**Table 2.**
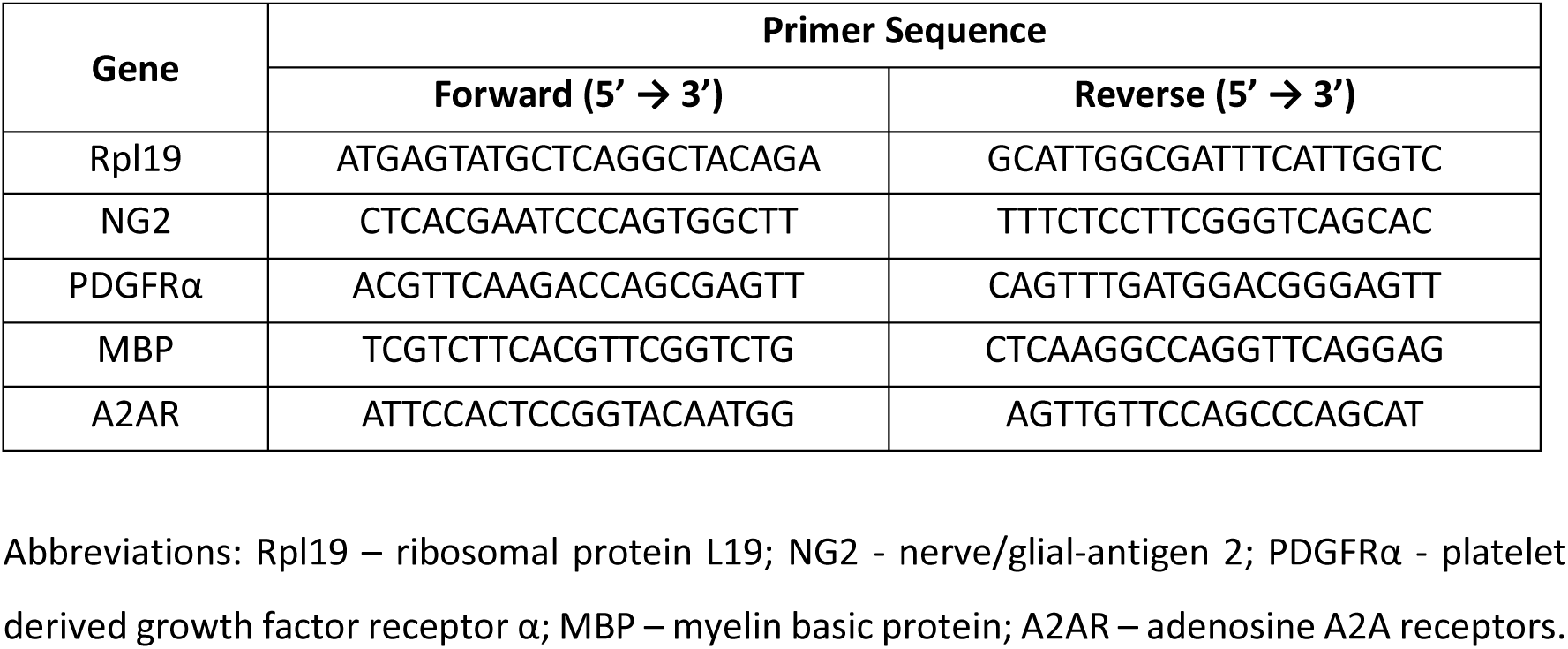
Primer Sequences.

### 6. Cell Viability Study

To study the effect of the pharmacological treatments on cell viability, we used propidium iodide (PI), a fluorescent DNA-binding dye widely employed to distinguish between healthy and apoptotic nuclei by binding to the DNA of cells with compromised cell membrane[21]. For this, SVZ cells were exposed at DIV 1 to 3 µg/mL of PI (#P4170, Sigma-Aldrich) for 30 min (**Suppl. Fig. 1A)**. Cells were then fixed in 4% paraformaldehyde (PFA) for 30 min and washed with phosphate buffered saline (PBS) at room temperature (RT). Nuclei counterstaining was performed with Hoechst 33342 (12 µg/mL #H1399, Thermo Fisher Scientific) for 30 min and after coverslips were mounted in Mowiol fluorescent medium (#324590, Sigma-Aldrich).

### 7. Cell Proliferation Study

To evaluate cell proliferation, we used 5-bromo-2’-deoxyuridine (BrdU) (#B5002, Sigma-Aldrich), a thymidine analogue able to substitute thymidine in the DNA double chain synthesis occurring in proliferating cells (Kee *et al*., 2002). Thus, SVZ cells at DIV 1 were exposed to 10 µM BrdU for the last 4 h of each pharmacological treatment (**Suppl. Fig. 1A)**. Then, cells were fixed in 4% PFA for 30 min and rinsed with PBS at RT. Afterwards, BrdU was unmasked by permeabilizing cells in PBS 1% Triton^TM^ X-100 (#X100, Sigma-Aldrich) at RT for 30 min and DNA was denatured in 1 M HCl for 20 min at 37°C. Following a 1.5 h incubation in PBS with 0.5% Triton^TM^ X-100 and 3% bovine serum albumin (BSA) to block nonspecific binding sites, cells were incubated overnight (O/N) at 4°C with the anti-mouse BrdU antibody (Table 2). After rinsing with PBS, cells were incubated for 2 h at RT in the corresponding secondary antibody (Table 3) and nuclei counterstaining with Hoechst 33342 in PBS, as described above. The final preparations were mounted using Mowiol fluorescent medium.

**Table 3.** Antibodies used.

| Primary Antibodies |  |  |  |  |
| --- | --- | --- | --- | --- |
| Antigen | Host | Dilution | Cat. Number | Company |
| BrdU | Mouse | 1:500 | #339802 | Biolegend |
| NG2 | Rabbit | 1:200 | #B5320 | Merck Milipore |
| PDGFRα<br>(CD140a) | Rat | 1:500 | #558774 | BD |
| MBP | Rabbit | 1:200 | #78896S | Cell Signalling |
| Secondary Antibodies |  |  |  |  |
| Antigen | Host | Dilution | Cat. Number | Company |
| Alexa Fluor<sup>®</sup> 488<br>anti-mouse | Donkey | 1:500 | #A21202 | Life Technologies |
| Alexa Fluor<sup>®</sup> 568<br>anti-rabbit | Donkey | 1:500 | #A10042 |  |
| Alexa Fluor<sup>®</sup> 488<br>anti-rat | Donkey | 1:500 | #A21208 |  |

Abbreviations: BrdU - 5-bromo-2'-deoxyuridine; NG2 - nerve/glial-antigen 2; PDGFRα - platelet derived growth factor receptor $\alpha$ ; MBP – myelin basic protein.

### 8. Cell Differentiation Study

SVZ-neurosphere derived cells maintained for 2 (**Fig. 1A**), 7 (**Fig. 2A**), and 14 (**Fig. 3A**) DIV with the pharmacological treatments were fixed for 30 min in 4% PFA in PBS, permeabilized and blocked for non-specific binding sites for 1.5 h in 0.5% Triton^TM^ X-100 and 3% BSA in PBS. Cells were then incubated O/N at 4°C with a combination of anti-NG2 and anti-PDGFRα (Table 3) for the labelling of OPCs or with anti-MBP (Table 3) to label mature OLs in 0.1% Triton X-100 and 0.3% BSA in PBS. The following day, cells were incubated for 2 h in the appropriate secondary antibody (Table 3) and nuclei counterstaining with Hoechst 33342 in PBS (see above). Coverslips were then mounted in Mowiol fluorescent medium.

**Fig. 1.**
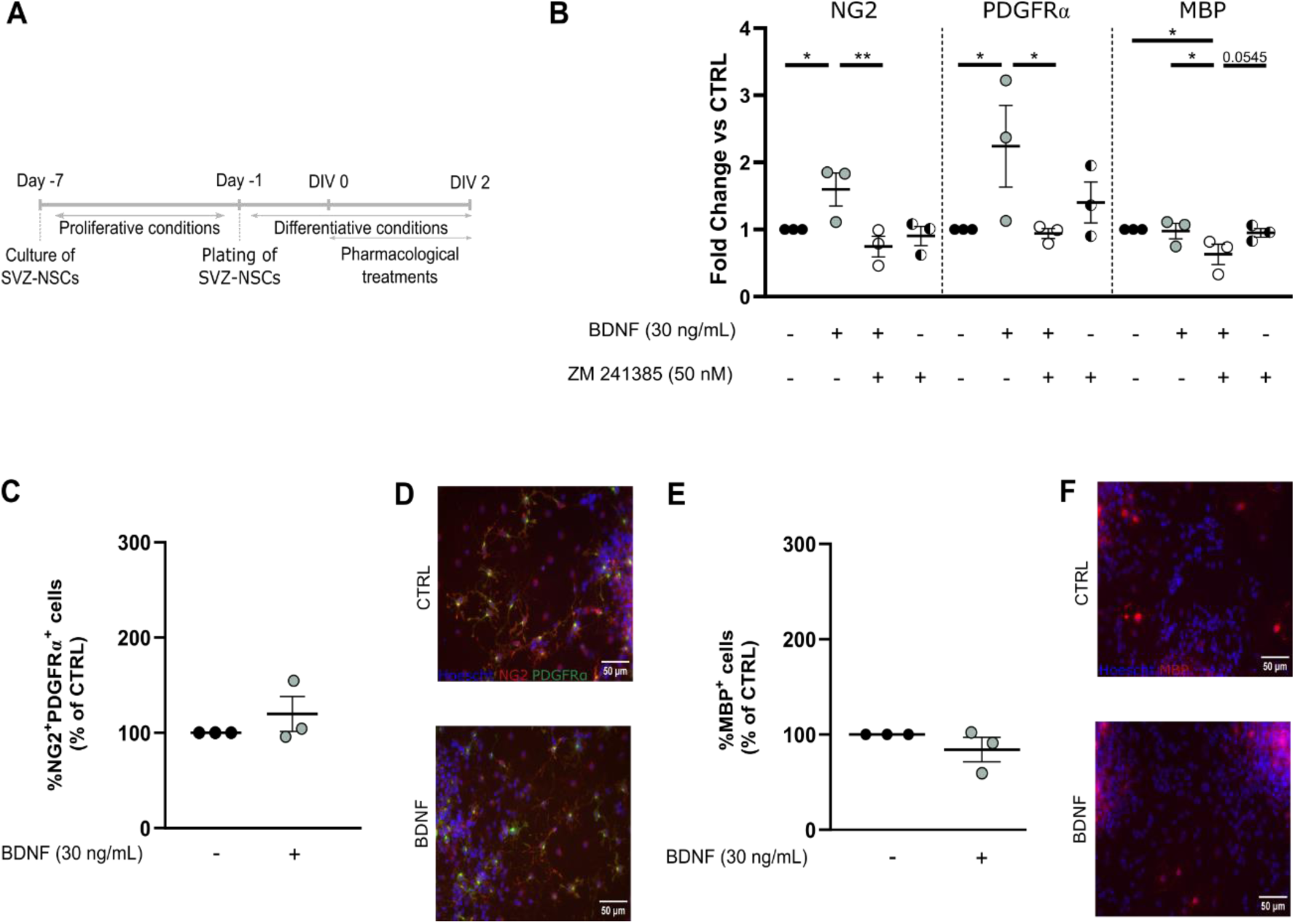
In SVZ neurospheres, BDNF induces an increase in the mRNA expression levels of the OPC markers NG2 and PDGFRα at DIV2, while the percentage of NG2^+^PDGFRα^+^ cells remains unchanged. Schematic representation of the experimental protocol. Cultures were performed at day -7, neurospheres plated for 24 h on day -1 and then cells were exposed to the pharmacological treatments for 2 days (DIV 2) (**A**). Graph depicts the mRNA expression levels of the oligodendroglial markers NG2, PDGFRα and MBP at DIV 2 (**B**). The graphs illustrate the percentage of NG2^+^PDGFRα^+^ (**C**) and MBP+ (**E**) cells derived from SVZ-NSCs at DIV 2. Representative images for OPCs from SVZ-NSCs stained for NG2 (red) and PDGFRα (green) (**D**), and OLs stained for MBP (red) (**F**) and counterstained for cell nuclei with Hoechst 33342 (blue). Values were normalized for the control mean for each experiment. Data presented as mean±SEM. One-way ANOVA, uncorrected Fisher’s LSD (*p<0.05; **p<0.01). n=3 independent cultures. Scale bar=50 µm. Abbreviations: SVZ-NSCs – SVZ neural stem cells; DIV – days in vitro; CTRL – control; BDNF – Brain-derived neurotrophic factor; NG2 - nerve/glial-antigen 2; PDGFRα - platelet derived growth factor receptor α; MBP – myelin basic protein

**Fig. 2.**
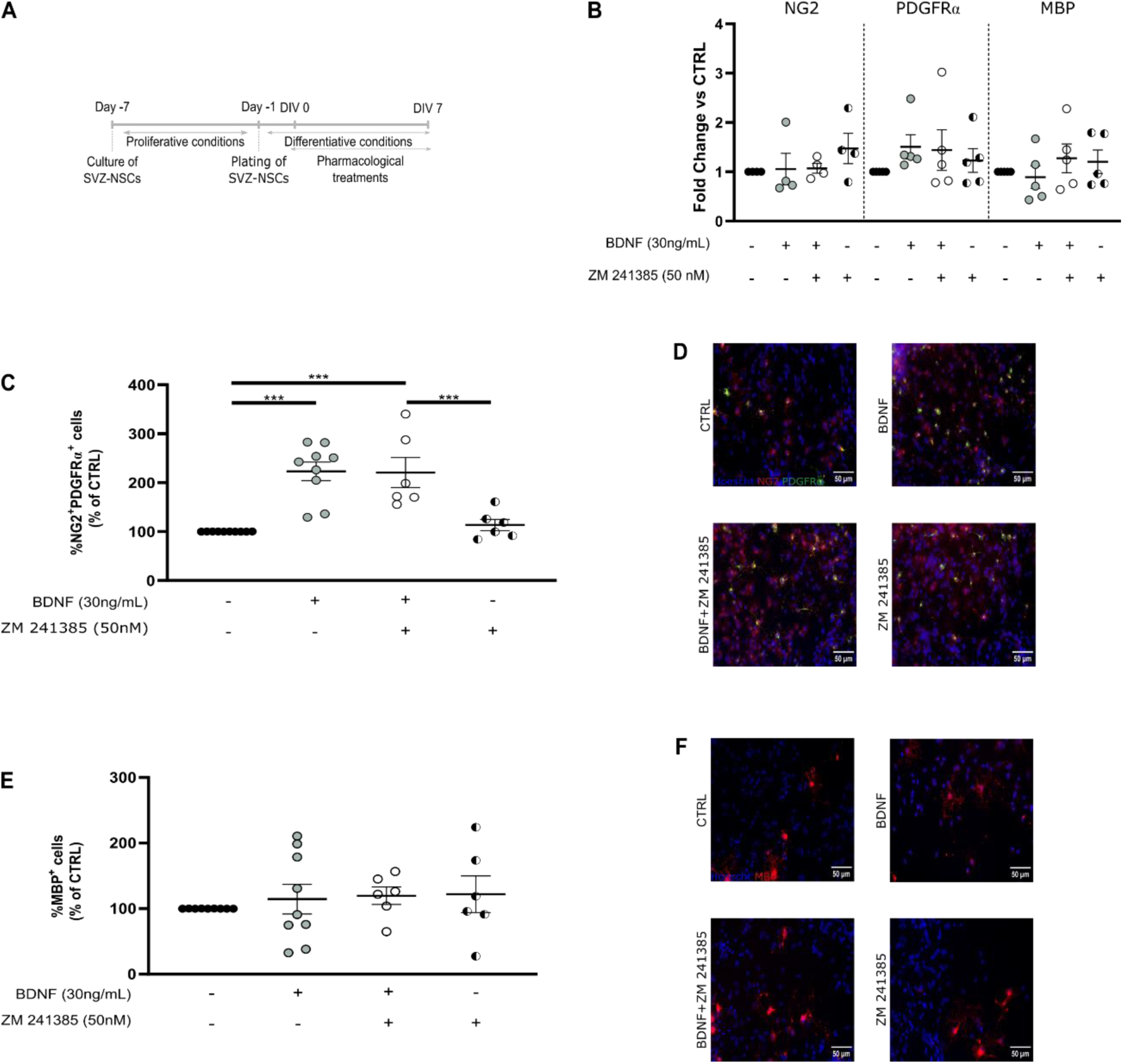
BDNF increases the differentiation of SVZ-NSCs into OPCs at DIV 7. Schematic representation of the experimental protocol. Cultures were performed at day -7, neurospheres plated at day -1 for 24 h and then cells were exposed to pharmacological treatments for 7 days (DIV 7) (**A**). The graph depicts the mRNA expression levels of the oligodendroglial markers NG2, PDGFRα and MBP at DIV 7 (**B**). The graphs illustrate the percentage of NG2^+^PDGFRα^+^ (**C**) and MBP+ (**E**) cells derived from SVZ-NSCs at DIV 7. Values were normalized for the control mean for each experiment. Representative images for OPCs from SVZ-NSCs, stained for NG2 (red) and PDGFRα (green) (**D**), and OLs, stained for MBP (red) (**F**) and counterstained for cell nuclei with Hoechst 33342 (blue). Data presented as mean±SEM. One-way ANOVA, uncorrected Fisher’s LSD (***p<0.001). n=4-10 independent cultures. Scale bar=50 µm. Abbreviations: SVZ-NSCs – SVZ neural stem cells; DIV – days in vitro; CTRL – control; BDNF – Brain-derived neurotrophic factor; NG2 - nerve/glial-antigen 2; PDGFRα - platelet derived growth factor receptor α; MBP – myelin basic protein

**Fig. 3.**
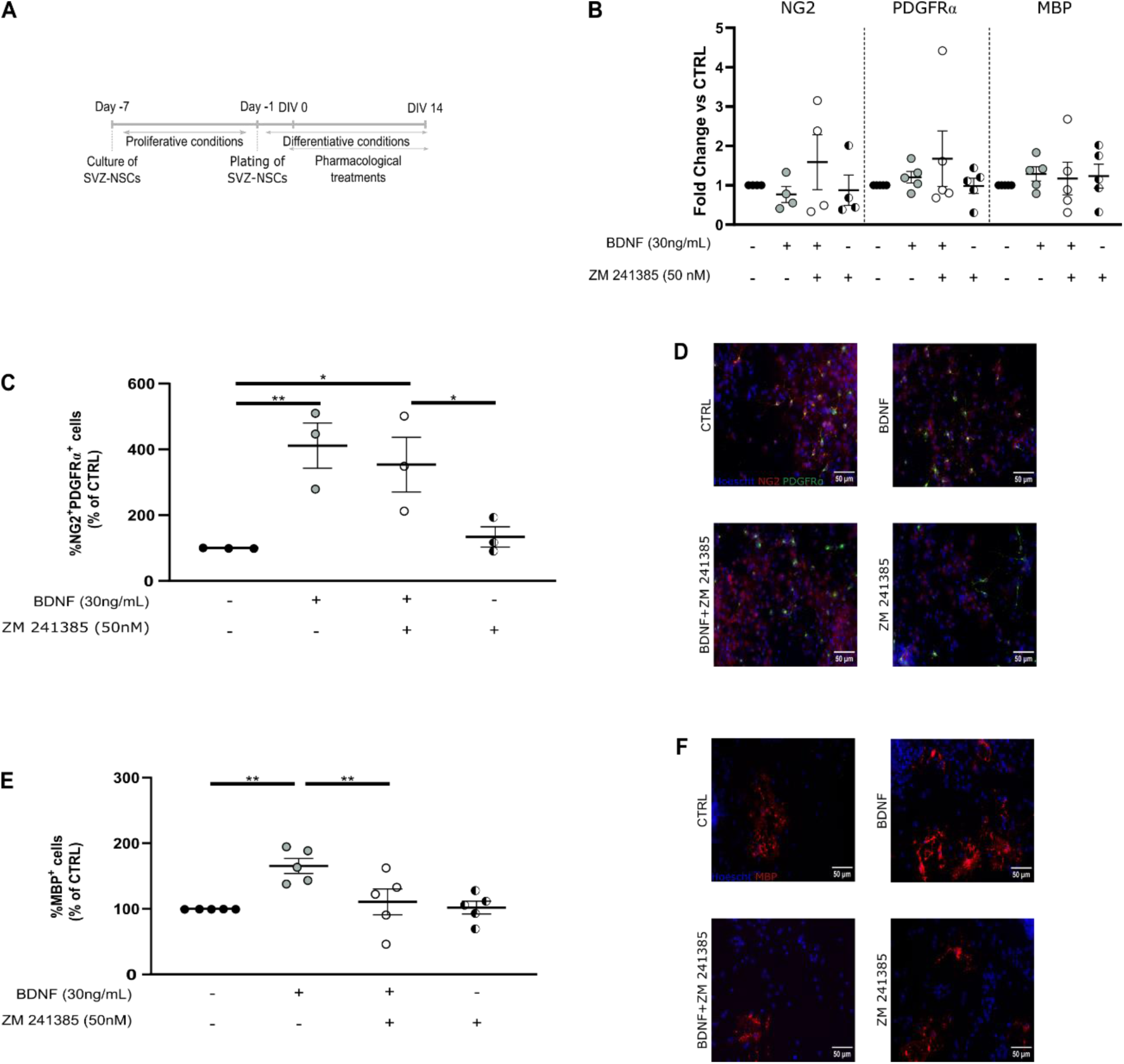
BDNF promotes the differentiation of SVZ-NSCs into OLs at DIV 14, an effect blocked by an A2AR antagonist. Schematic representation of the experimental protocol. Cultures were performed at day -7, neurospheres plated at day -1 for 24 h and then cells were exposed to pharmacological treatments for 14 days (DIV 14) (**A**). The graphs depict the mRNA levels of oligodendroglial markers NG2, PDGFRα and MBP (**B**) and the percentage of NG2^+^PDGFRα^+^ (**C**) and MBP+ (**E**) cells derived from SVZ-NSCs at DIV 14. Values were normalized for the control mean for each experiment. Representative images for OPCs from SVZ-NSCs, stained for NG2 (red) and PDGFRα (green) (**D**) and OLs, stained for MBP (red) (**F**) and counterstained for cell nuclei with Hoechst 33342 (blue). Data presented as mean±SEM. One-way ANOVA, uncorrected Fisher’s LSD (*p<0.05; **p<0.01). n=3-5 independent cultures. Scale bar=50 µm. Abbreviations: SVZ-NSCs – SVZ neural stem cells; DIV – days in vitro; CTRL – control; BDNF – Brain-derived neurotrophic factor; NG2 - nerve/glial-antigen 2; PDGFRα - platelet derived growth factor receptor α; MBP – myelin basic protein

### 9. Morphological analysis

Individualized MBP+ OLs were used to perform morphology analysis at DIV 7 and DIV 14, using the Neuroanatomy plugin from ImageJ software (http://fiji.sc/). Sholl analysis was performed by drawing concentric circles around the cell body with an increasing radius of 5 μm.

### 10. Microscopy

Fluorescence images were captured using an AxioCamMR3 monochrome digital camera (Carl Zeiss Inc.) mounted on an Axiovert 200 inverted widefield fluorescence microscope (Carl Zeiss Inc.), with a 40x objective. Images were recorded using the software AxioVision 4 (Carl Zeiss Inc.). The pixel size in the object space was 0.25 μm and the captured image size was 1388 × 1040 pixels. Images were stored and analysed in an uncompressed 8-bit Tiff format. Cells were counted manually using ImageJ software.

### 11. Statistical Analysis

In the immunocytochemistry experiments, images were acquired at the border of SVZ neurospheres, where migrating cells form a pseudo-monolayer of cells. Every independent experiment (n) corresponds to one independent SVZ neurosphere culture obtained from one litter of C57BL/6J mice at P1-P3. In every independent experiment, each condition was analysed in triplicate, i.e., using three different coverslips. In each coverslip, the percentages of labelled cells were calculated from cell counts in five independent microscopic fields, with a 40x objective (approximately 100-250 cells per field). Therefore, each n value corresponds at least to the average of 3×5×100 cells. In each set of experiments, data obtained were normalized to each corresponding control. Data are expressed as mean± standard error of the mean (SEM) and the control was set to 100%.

Regarding morphological analysis, for each condition 12-25 cells were analysed (per independent culture). For Sholl analysis plots, mean values for the total number of cells per condition were taken and are expressed as mean±SEM. Regarding OL area, maximum intersections and distance from soma with maximum intersections are represented as data with median. For the area graphical representations, control was set to 100%.

For the qRT-PCR, SVZ-derived cells were also grown in triplicates and each independent culture was considered n = 1. Values were normalized to the control expression of Rpl19 expression for each experiment. Data presented as mean ± SEM and the control was set to 1.

All statistical analyses were performed using GraphPad Prism version 8 for windows (GraphPad Software, Inc). Statistical significance was determined using Two-way ANOVA or One-way analysis, when appropriate, followed by uncorrected Fisher’s LSD test, with p<0.05 considered to represent statistical significance. Values outside the mean±2SD (standard deviation) range were classified as significant outliers and excluded from further analyses[22, 23].

All artwork was designed by the author using Inkscape (http://www.inkscape.org/).

## Results

### BDNF stimulates the mRNA expression levels for OPC markers at DIV 2, an effect blocked by an A2AR antagonist

To follow the differentiation process from precursors to myelinating oligodendrocytes, SVZ-derived neurospheres were treated with BDNF and the A2AR antagonist ZM 241385 under differentiative conditions for 1, 2, 7 or 14 days. At DIV 1 **(Suppl. Fig. 1A)**, neither BDNF (30 ng/mL) nor ZM 241385 (50 nM) significantly impacted cell survival (**Suppl. Fig. 1B, C**) nor proliferation (**Suppl. Fig. 1D, E**).

Importantly, co-treatment with BDNF (30 ng/mL) and ZM 241385 (50 nM) led to a significant decrease in A2AR mRNA expression levels when compared with BDNF (0.650-fold relative to CTRL; n=3; p<0.05), while ZM 241385 (50 nM) alone reduced A2AR mRNA expression levels when compared with control (0.383-fold relative to CTRL; n=3; p<0.05) (**Suppl. Fig. 2).** For the later timepoints, DIV 7 and DIV 14, no significant effects were observed.

Regarding oligodendrocyte differentiation at DIV 2, SVZ cells treated with BDNF (30 ng/mL) (**Fig. 1A**) showed a significant increase in the mRNA expression levels for the OPC markers, NG2 and PDGFRα (NG2: 1.597-fold and PDGFRα: 2.240-fold relative to CTRL; n=3; p<0.05). Interestingly, this effect mediated by BDNF was blocked when SVZ cells were co-treated with the A2AR antagonist, ZM 241385 (50 nM) (**Fig. 1B**). For MBP, a marker for mature myelinating OLs, a decrease in the expression levels was observed when cells were treated with both BDNF and ZM 241385 (0.6300-fold relative to CTRL; n=3; p<0.05), while each treatment alone was virtually devoid of effect (**Fig. 1B**). No changes in the percentage of NG2^+^PDGFRα^+^ OPCs (**Fig. 1C, D**) nor MBP^+^ mature OLs (**Fig. 1E, F**) derived from SVZ-NSCs were observed in this timepoint after treatment with BDNF.

Overall, these results indicate that, at DIV 2, BDNF and A2AR affect OPCs but not OLs, and effects are only evident at the mRNA level.

### BDNF treatment for 7 DIV, promotes the differentiation of SVZ-NSCs into OPCs

BDNF (30 ng/mL) treatment for 7 DIV (**Fig. 2A**) did not induce changes in the mRNA expression levels for NG2, PDGFRα and MBP (n=4-5, p>0.05) (**Fig. 2B**). Interestingly, our data shows that treatment with BDNF increased the percentage of NG2^+^PDGFRα^+^ OPCs (CTRL: 100.0±3.378; BDNF: 220.8±16.58; n=9-10; p<0.001). The co-treatment with the A2AR antagonist had no effect on this potentiation, and the antagonist A2AR by itself also did not change the percentage of NG2^+^PDGFRα^+^ cells when comparing with control (**Fig. 2C, D**). Concerning mature OLs, no changes were observed in the percentage of MBP^+^ cells derived from SVZ-NSCs (n=6-9; p>0.05) (**Fig. 2E, F**).

These data emphasize the role of BDNF in the differentiation into OPCs derived from SVZ-NSCs.

### BDNF treatment for 14 DIV, stimulates the differentiation of SVZ-NSCs into mature OLs, an effect blocked by an A2AR antagonist

SVZ-NSCs were maintained in culture for 14 days under differentiative conditions to fully accompany the process of differentiation into mature OLs (**Fig. 3A**). No changes were observed for the mRNA expression levels at DIV 14 (n=4-5, p>0.05) (**Fig. 3B**). Importantly, the BDNF effects detected while quantifying NG2^+^PDGFRα^+^ OPCs at DIV 7 were also evident at DIV 14 (CTRL: 100.0±14.90; BDNF: 399.1±46.63; n=3; p<0.01) (**Fig. 3C, D)**. Also, and as observed at DIV 7, the co-treatment with the A2AR antagonist had no effect on this potentiation, and treatment solely with the A2AR antagonist also did not affect the percentage of NG2^+^PDGFRα^+^ OPCs (**Fig. 3C, D)**. Importantly, however, this more prolonged treatment with BDNF led to an increase in the percentage of MBP^+^ mature OLs (CTRL: 100.0±0.009; BDNF: 165.3±11.49; n=5; p<0.01), which was blocked by co-treatment with ZM 241385 (BDNF+ZM 241385: 110.6±19.82; n=5; p<0.01 BDNF *vs* BDNF+ZM) (**Fig. 3E, F**).

Consistent with the previous results at earlier timepoints, these data suggest that BDNF supports the differentiation from OPCs to mature OLs, an effect blocked by an A2AR antagonist specifically in mature OLs.

### BDNF has distinct effects on OL morphology at DIV 7 and DIV 14

BDNF, when incubated for 7 DIV, significantly reduced the branching of MBP^+^ cells, as the distance from the soma increased, particularly between 40 – 80 µm, an effect that was not blocked by co-treatment with ZM 241385 (**Fig. 4A, E**). Consistent with this observation, the distance from the soma with the maximum number of branch points was significantly decreased in cells incubated with BDNF, and the co-treatment with ZM 241385 also did not change this effect (CTRL: 35.19±2.448; BDNF: 29.04±1.933; BDNF+ZM 241385: 28.98±2.107; n=52-59 cells; p<0.05) (**Fig. 4B**). No changes were observed in the maximum number of intersections (n=52-59 cells, p>0.05) (**Fig. 4C**), nor in the total area of the cells (**Fig. 4D**).

**Fig. 4.**
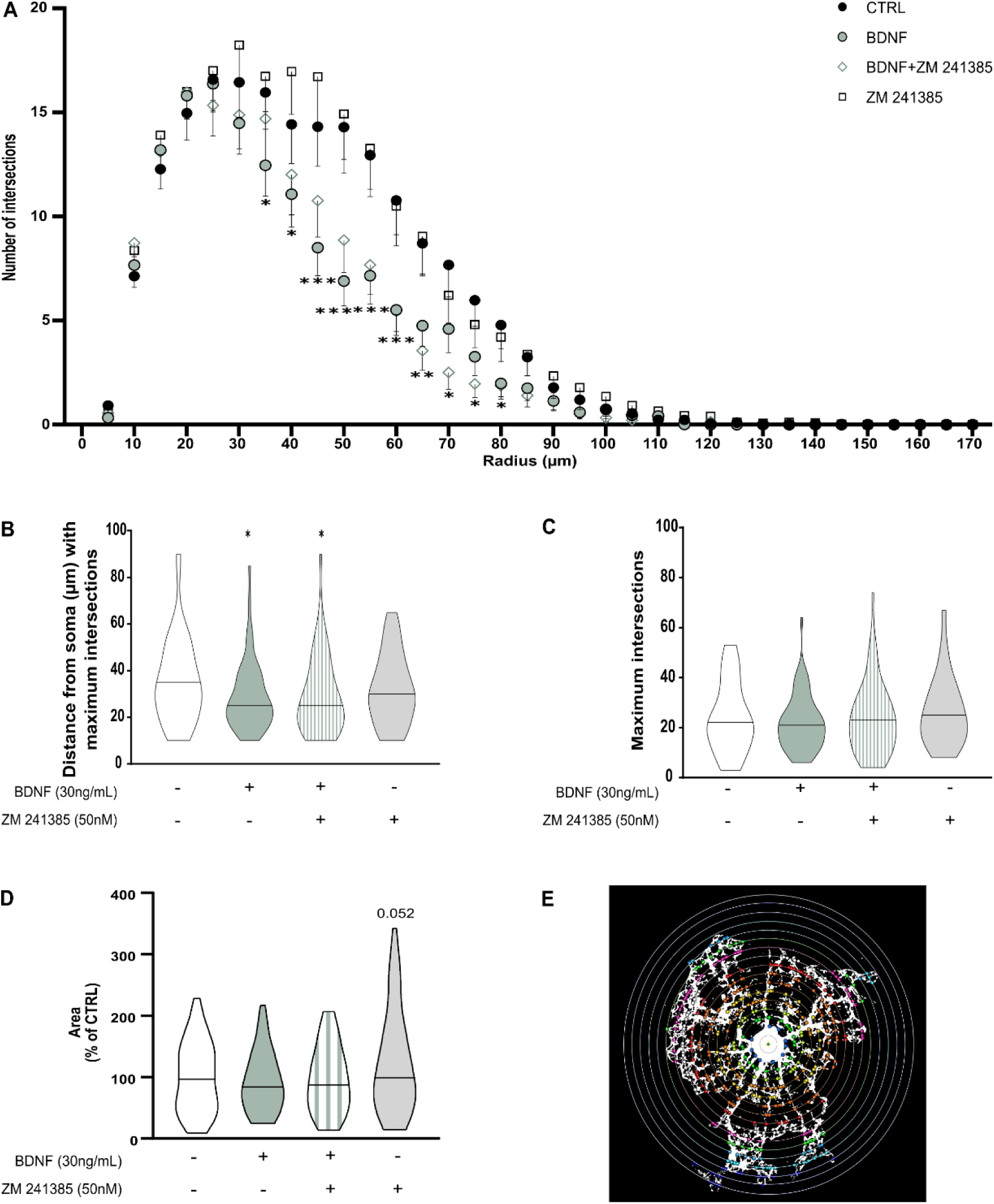
BDNF, at DIV7, decreases the ramified morphology of MBP^+^ OLs distal to the soma, an effect that is not blocked by the A2AR antagonist. Graph depicts the number of processes that intersect with the concentric circles of the Sholl analysis, as a function of the radius, i.e., the distance from the soma (**A**). The graphs depict distance from the soma with the maximum number of intersections (**B**), the maximum number of intersections (**C**), and the area (**D**). Sholl analysis was performed by placing a series of concentric circles spaced at 5 μm intervals centered on the soma (**E**). Data presented as mean±SEM (A) or data with median (B, C, D). Two-way ANOVA or One-way ANOVA, uncorrected Fisher’s LSD [(A): BDNF vs CTRL: ***p<0.001, **p<0.01, *p<0.05; BDNF vs BDNF + ZM 241385: p>0.05; (B): BDNF and BDNF+ZM 241385 vs CTRL: *p<0.05]. n=52-59 cells from 3 independent cultures. Abbreviations: CTRL – control; BDNF – Brain-derived neurotrophic factor; MBP – myelin basic protein

Notably, at DIV 14, the effect of BDNF upon branching was no longer evident since there were no differences in the number of intersections of the OLs for all treatment conditions (**Fig. 5A, E**). No changes were observed in the distance from the soma with the maximum number of branch points (**Fig. 5B**), nor in the maximum number of intersections (**Fig. 5C**) (n=45-48 cells; p>0.05). Interestingly, the area of the cells was significantly increased after treatment with BDNF, an effect not blocked with the co-treatment with ZM 241385 (CTRL: 100.0±48.85; BDNF: 121.6±56.11; BDNF+ZM 241385: 121.5±60.22; n=45-48 cells; p<0.05) (**Fig. 5D**).

**Fig. 5.**
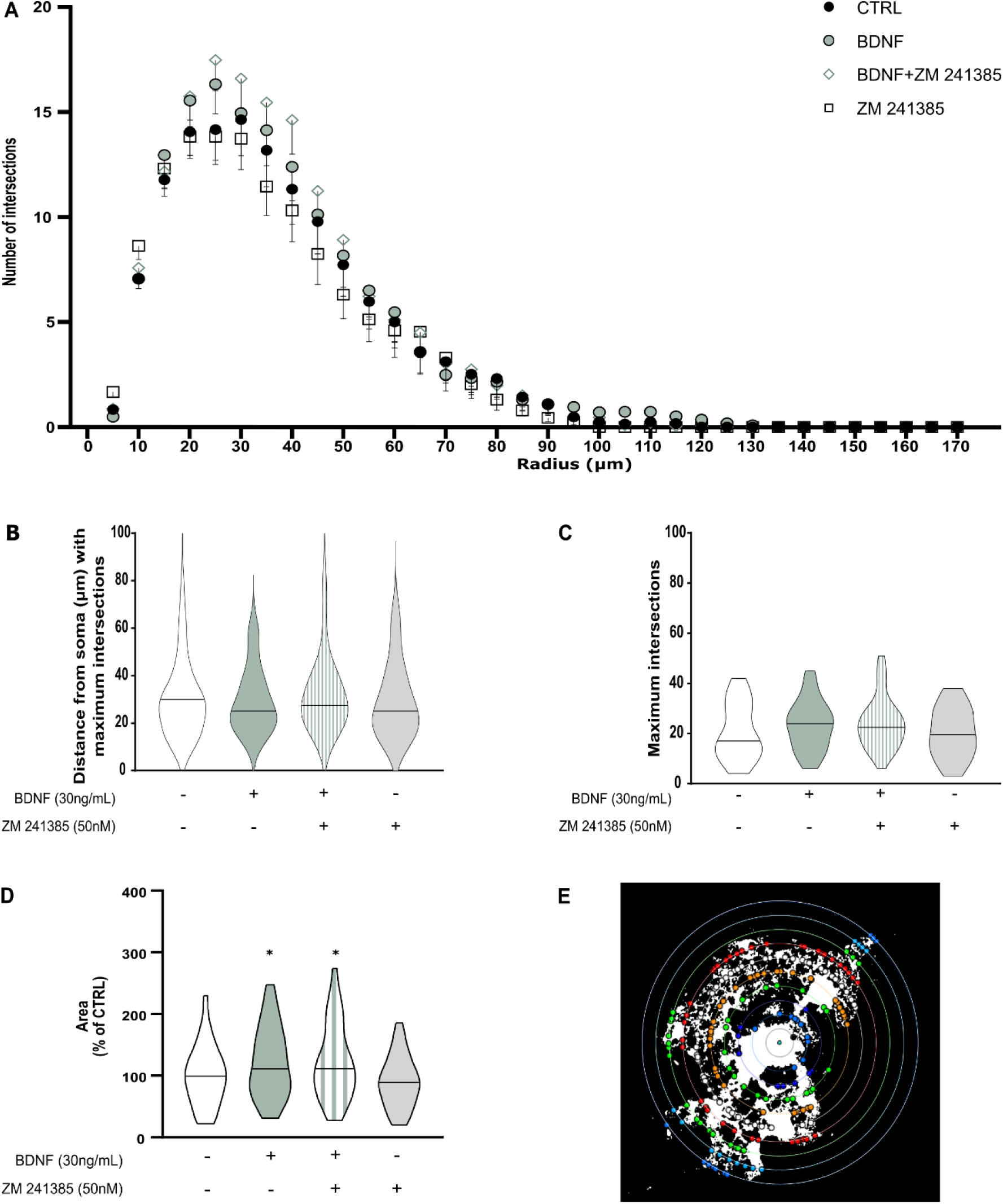
BDNF, at DIV 14, has no effect on the morphology of MBP^+^ cells. Graph depicts the number of processes that intersect with the concentric circles of the Sholl analysis, as a function of the radius, i.e., the distance from the soma (**A**). The graphs depict distance from the soma with the maximum number of intersections (**B**), the maximum number of intersections (**C**), and the area (**D**). Sholl analysis was performed by placing a series of concentric circles spaced at 5 μm intervals centered on the soma (**E**). Data presented as mean±SEM (A) or data with median (B, C, D). Two-way ANOVA or One-way ANOVA, uncorrected Fisher’s LSD [(D): BDNF and BDNF+ZM 241385 vs CTRL: *p<0.05)]. n=45-48 cells from 3 independent cultures. Abbreviations: CTRL – control; BDNF – Brain-derived neurotrophic factor; MBP – myelin basic protein

Altogether, these data show that BDNF exposure for 7 DIV reduced the complexity and extent of less mature cells, while at DIV 14 there were no morphological differences in BDNF exposed/non exposed cells, suggesting that the effect mediated by BDNF is limited to the OL maturation process.

## Discussion

BDNF plays a well-documented role in the formation and maturation of OLs, making it a critical target for addressing demyelinating diseases[9, 13, 24]. The interaction between BDNF and adenosine A2ARs has previously been implicated in regulating synaptic transmission[18, 25, 26] and postnatal neurogenesis[20]. As summarized in **Fig. 6**, a main finding of the present work is that the influence of BDNF upon OL-derived from SVZ neurospheres is time-dependent, promoting the expression of OPC mRNA markers on an initial stage, followed by enhanced OPC and OL cellular differentiation at later stages. Moreover, BDNF induces morphological changes in OL with distinct effects at different timepoints. Importantly, the effects observed in the percentage of MBP^+^ cells in late differentiation for mature OLs were dependent of active A2AR. In our study, both BDNF and the A2AR antagonist had no effect on cell survival nor proliferation. Previous work from our group performed in rat showed that treatment with BDNF and the A2AR antagonist does not affect cell viability, however BDNF significantly increase cell proliferation in neurospheres from the hippocampal dentate gyrus[20]. This discrepancy can be explained by the different neurogenic niche being studied, and/or species used. Other groups have studied how BDNF can modulate SVZ-NSCs and showed that intraventricular administration of BDNF results in decreased proliferation rate in the SVZ of mice[27]. However, others have shown that BDNF has no effects on the proliferation of cultured spinal cord NSCs[28]. Moreover, an anti-apoptotic effect of BDNF has been described in a model of murine NSCs, an effect mediated by TrkB receptor activation after neurotoxin-induced apoptotic cell death[29]. The role of purinergic receptors in the differentiation of stem cells has been widely studied[30]. Regarding A2ARs, its blockade during the process of embryogenesis does not affect SVZ proliferation[31]. Notably, Stafford *et al.* previously showed that activation of A2AR leads to a decrease in cell proliferation of SVZ-primary neurospheres, likely initiating neuronal differentiation[32]. Our group has demonstrated that A2AR activation protects rat DG-derived immature neurons from cell death contributing to their differentiation into the neuronal lineage. Furthermore, we also showed that the activation of A2AR receptors is essential for the beneficial effects of BDNF on neuronal differentiation[20].

**Fig. 6.**
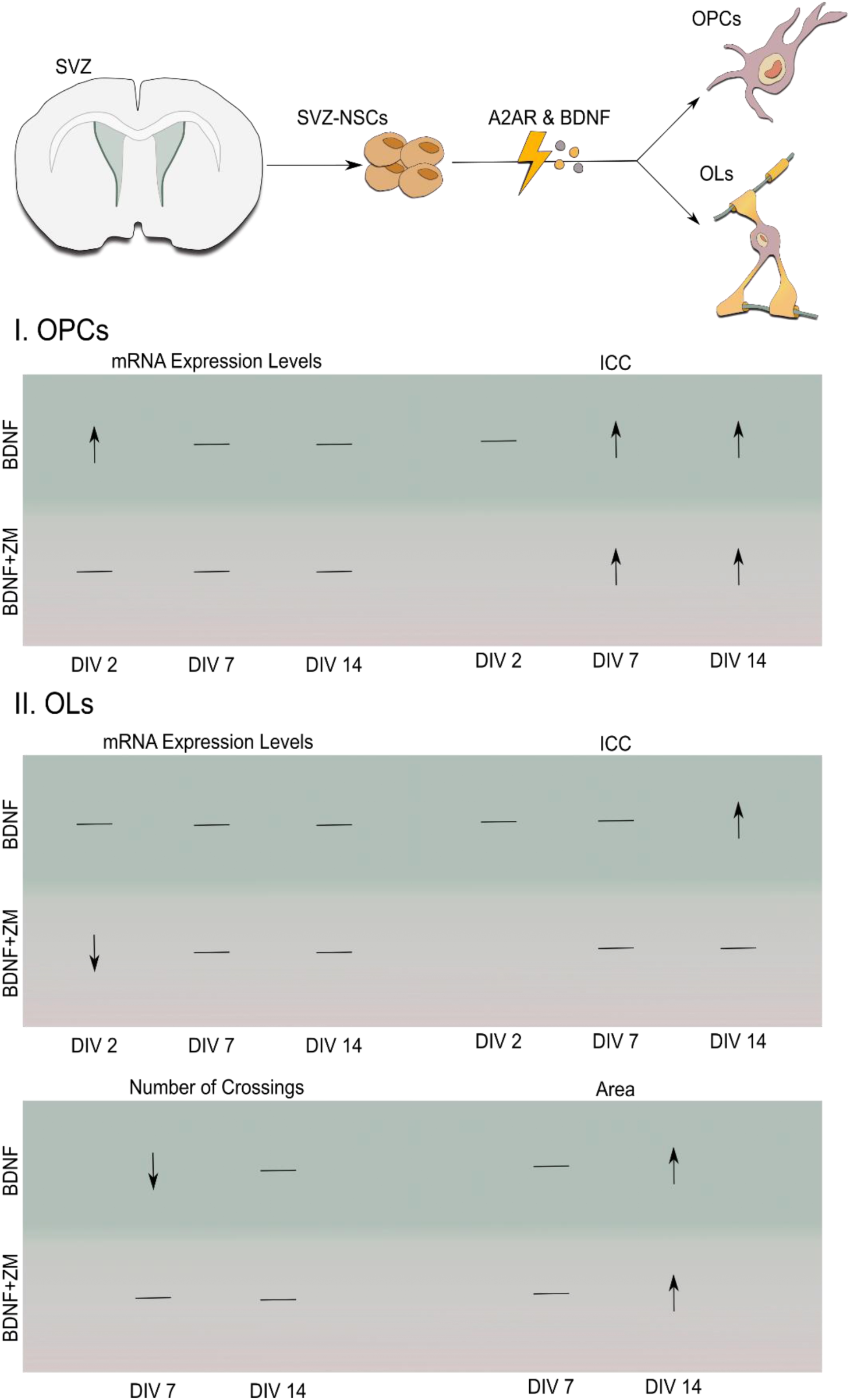
This work was focused on the modulation of SVZ-derived NSCs by BDNF and an A2AR antagonist (ZM 241385), and how these treatments affected differentiation into the oligodendroglial lineage. Regarding OPCs, BDNF increased mRNA levels for markers of these precursors at DIV 2, whereas the effects on protein expression observed using ICC were only observed at DIV 7 and DIV 14. At both timepoints, A2AR antagonist did not block the modulatory effects of BDNF, maintaining the increased percentage of OPCs. Regarding mature OLs, treatment with BDNF had no effect, however co-treatment with the A2AR antagonist led to a decrease in mRNA expression levels for MBP, a marker associated with myelinating OLs. Effects on ICC were only seen at DIV 14, where BDNF led to an increase in the percentage of MBP^+^ cells. Regarding the morphology of OLs, results showed that at DIV 7 there were no changes in area of the cells, but there was a decrease in branching of BDNF-treated OLs. Interestingly, at DIV 14 the effects observed in the previous timepoint with BDNF treatment were lost, and all treatment conditions promoted differentiation of MBP^+^ cells with similar morphology. Nevertheless, at DIV 14 BDNF promoted an increase in the area of MBP^+^ cells, an effect not blocked by the A2AR antagonist. SVZ – Subventricular Zone; SVZ-NSCs – SVZ-derived neural stem cells; A2AR – Adenosine A2A receptor; BDNF – brain-derived neurotrophic factor; DIV – days in vitro; OPC – oligodendrocyte precursor cell; OL – oligodendrocyte; ICC – immunocytochemistry; ZM – ZM 241385, A2AR antagonist. Arrows indicate increase or decrease for that pharmacological condition; horizontal bars indicate no change when comparing with control

To evaluate oligodendrocyte differentiation, we assessed the gene expression levels of oligodendroglial markers NG2, PDGFRα, and MBP at DIV 2, 7, and 14. Interestingly, changes in mRNA expression were observed only at DIV 2, where BDNF treatment led to increased levels of NG2 and PDGFRα mRNA. For OPC identification, we used a double-label immunocytochemistry strategy with NG2 and PDGFRα, confirming the commitment of these cells to the oligodendroglial lineage. BDNF treatment resulted in an increased percentage of NG2^+^PDGFRα^+^ OPCs at both DIV 7 and DIV 14, while the percentage of MBP^+^ mature OLs increased only at DIV 14. This pattern suggests that the initial upregulation of OPC markers by BDNF at DIV 2 may facilitate the differentiation into MBP^+^ OLs observed at DIV 14. Therefore, the effects observed showed temporal regulation, with primary effects on mRNA levels that only posteriorly were observed on protein expression. Several studies have approached the temporal evolution of mRNA to protein expression, namely going in depth to transcriptome and proteomic analysis. Cheng *et al*. used an endoplasmic reticulum stress model in mammalian cells to demonstrate that both mRNA and protein regulatory mechanisms are relevant, but whereas mRNA concentrations shift in a transient pattern, returning to pre-stress values, protein concentrations switch one time and remain at a new steady state[33]. The regulation of both mRNA and protein expression, and their time-dependency, are crucial targets for putative therapeutic interventions. Therefore, the identification of molecules and their receptors that may modulate them is of high importance. Work by Langhnoja *et al*. [24], using in vitro cortical NSC cultures, also showed that BDNF potentiates the differentiation of NSCs into the oligodendroglial lineage, an observation compatible with our findings. While they have shown that BDNF promotes NSC differentiation into the oligodendroglial lineage in a dose-dependent manner [24], we could demonstrate that this occurs in a time-dependent manner. Interestingly, previous in vitro and in vivo works have shown that after a demyelinating stimulus, SVZ-NSCs differentiate into both OPCs and OLs that can migrate to the lesion areas [7, 34, 35]. Importantly, Vondran *et al*., using BDNF^+^/^-^ mice subjected to a cuprizone demyelination model, concluded that the levels of this neurotrophin impact in the proportion of OPCs, as well as upon their ability to maturate and produce myelin proteins[36].

BDNF also plays a role in shaping OL morphology. At DIV 7, BDNF led to a reduction in ramification complexity, an effect that was not blocked by the A2AR antagonist. However, at DIV 14, the effect of BDNF was no longer present, since OLs exhibited similar morphology across all treatment conditions. These results suggest that BDNF impacts cell morphology during the maturation process, but not in fully mature cells. OLs present distinct morphologies throughout their differentiation process and vary by brain region[37, 38]. Previous studies have documented structural changes in the OL cytoskeleton, from process elongation to myelin sheath production, in vitro [37, 39, 40]. Our observation of decreased branching of MBP^+^ OLs treated with BDNF at DIV 7, which was lost by DIV 14, may reflect a role of BDNF in steps related to cell growth and process elongation until the maturation process is complete. While our data revealed a role for BDNF during the OL maturation process, the mechanisms behind the effect observed, whether it is a signalling cascade resulting in activation or downregulation of other critical factors in this process, remains to be further assessed.

Markedly, our work highlights that OLs have a relatively long maturation process, that the action of BDNF mostly affects the later stages of their differentiation, where we observed a reduction in BDNF potentiation upon blocking A2ARs, highlighting the crosstalk between BDNF and adenosine A2AR during SVZ-derived oligodendroglial differentiation. Considering our timepoints and experimental conditions, our data suggest that BDNF-treated cells differentiate into mature myelinating OLs expressing MBP by DIV 14. However, in the presence of both BDNF and the A2AR antagonist, OL differentiation seems to be delayed. Numerous studies from our group and others have explored the A2AR-dependent activity of BDNF in hippocampal activity[41–43] and in animal models of diseases such as epilepsy[44], Huntington’s disease[45] and fragile X syndrome[46]. Also, other authors have described how blocking A2AR prevents the release of BDNF and impairs consequent microgliosis, an interaction restricted to pathological conditions, namely a lipopolysaccharide stimulus[47], in situations of motor neuron death [48], or age-related[49]. Our results align with previous findings, expanding the A2AR/BDNF interaction towards maturation of OLs from SVZ-derived OPCs, thus suggesting a possible impact upon remyelination under disease conditions. Also, while our data revealed a role for BDNF during the OL maturation process, the mechanisms behind this effect, whether it involves a signalling cascade leading to the activation or to downregulation of other critical factors, remains to be further investigated. Taken together, our data provide insights into the interactions between BDNF and A2AR signalling to modulate OL formation from SVZ-NSCs in vitro, under physiological conditions. These results provide relevant information to pave the way for future studies exploring BDNF and A2AR signalling as potential targets for promoting OL differentiation and remyelination under physiological and pathological conditions.

## Acknowledgments

The authors declare no conflict of interests. The authors would like to acknowledge the Rodent and Bioimaging facilities at Instituto de Medicina Molecular João Lobo Antunes (Lisbon, Portugal). We would like to acknowledge Dr. Joana Coelho (iMM, FMUL) that kindly provided the primers for A2ARs. This work was supported by Fundação para a Ciência e Tecnologia (FCT) fellowships PD/BD/150343/2019 and COVID/BD/153328/2023 (JMM) and 2020.04492.BD (JBM). This work has received funding by European Union’s Horizon Europe research and innovation programme under grant agreement n° 101160180 (PANERIS).

## Author Contributions

JMM conceived the study, designed and performed the experiments, analysed results and wrote the paper; BS and NP analysed results; AB and JBM collaborated in the experiments and participated in result discussion; AMS, AB, AF, SX participated in data interpretation and result discussion; AMS, AF, SX provided funding support and participated in the writing of the paper; SX designed and supervised the study.

## Data availability

The datasets generated during and/or analysed during the current study are not publicly available but are available from the corresponding author on reasonable request.

## Supplementary Figures

**Supplementary Figure 1.**
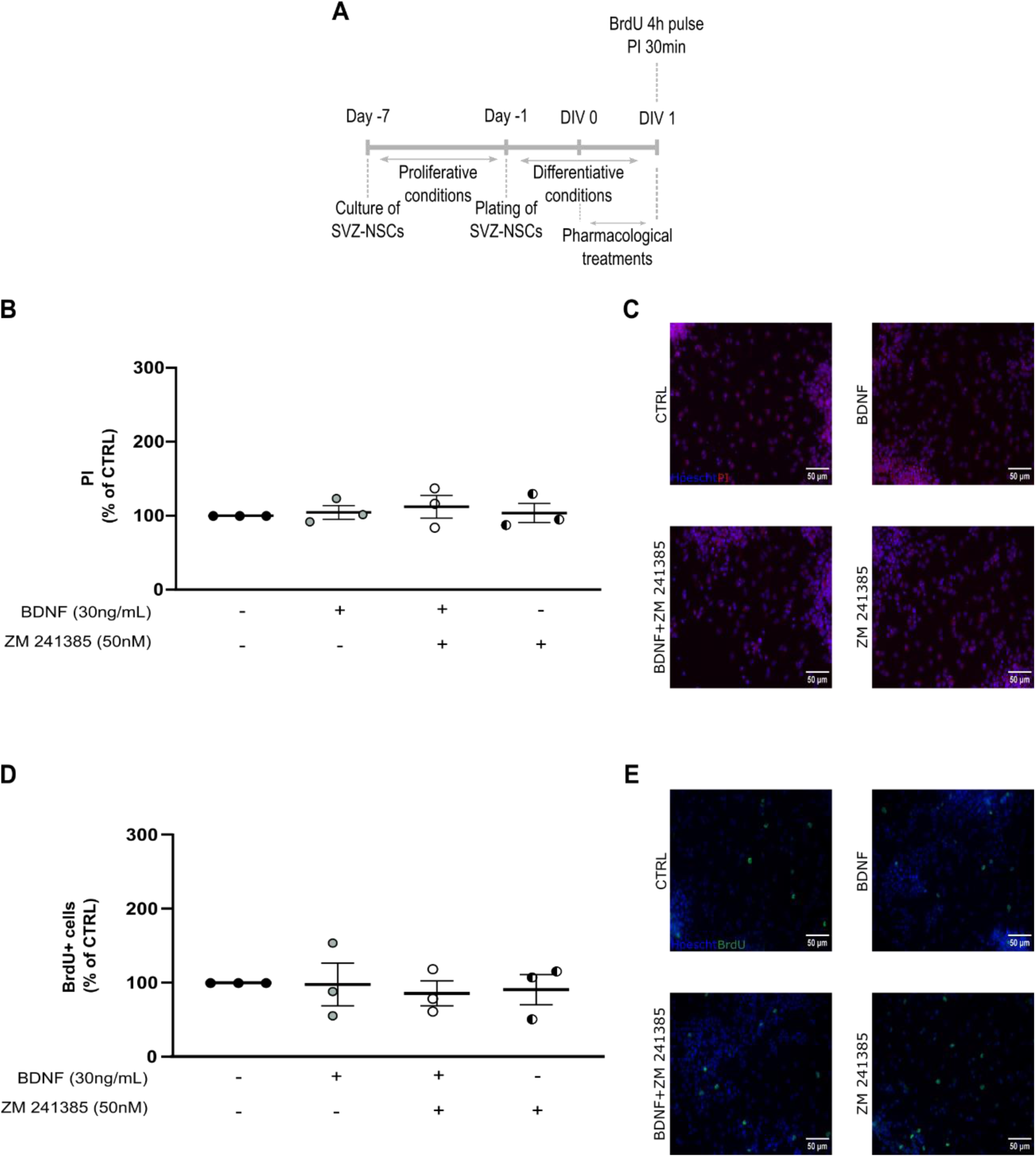
BDNF has no effect on cell survival nor proliferation at DIV 1. Schematic representation of the experimental protocol used to study cell proliferation and survival (**A**). Cultures were performed at day -7, neurospheres plated for 24h and then cells were exposed to pharmacological treatments for 24h (DIV 1). BrdU was incubated 4h and PI 30min before cell fixation. Graphs depict the percentage of PI^+^ cells (**B**) and BrdU^+^ cells (**D**) after the different treatments. Representative images for BrdU^+^ (**C)** and PI^+^ (**E**) cells stained for Hoechst 33342 (blue), BrdU (green) and PI (red), respectively. Scale bar=50 µm. Abbreviations: CTRL – control; BDNF – Brain-derived neurotrophic factor; PI – Propidium iodide, BrdU - 5-bromo-2’- deoxyuridine

**Supplementary figure 2.**
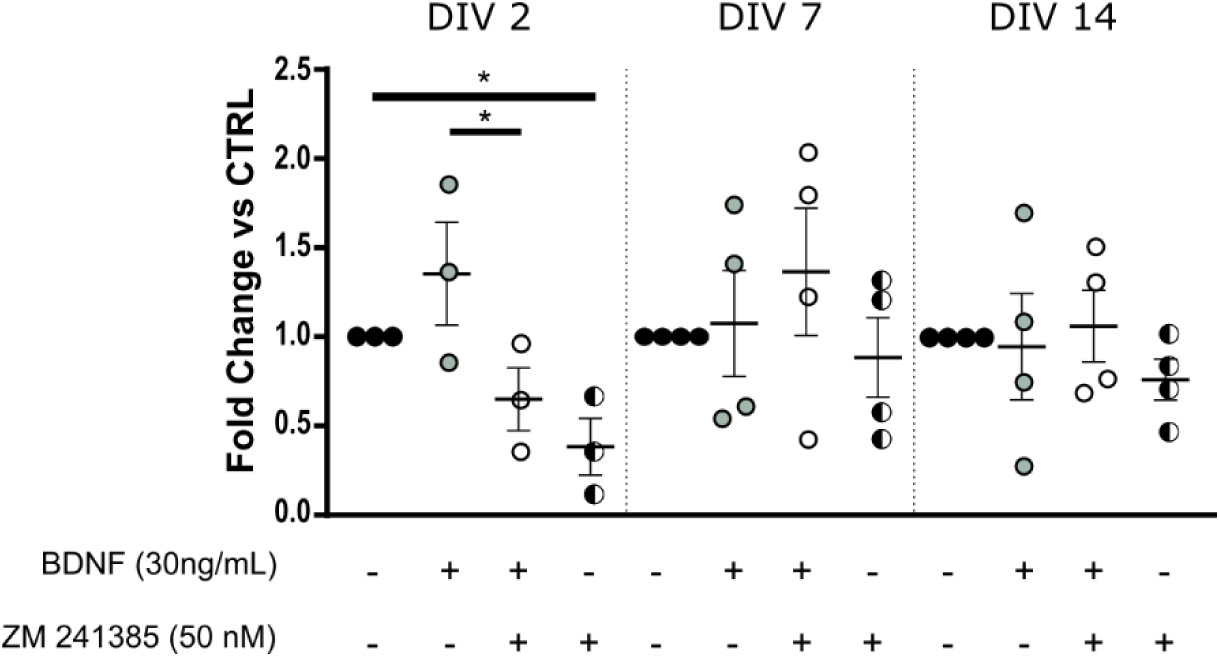
A2AR antagonist promotes a decrease in A2AR mRNA expression levels at DIV 2. Graph shows mRNA levels for the A2AR in the experimental timepoints tested: DIV 2, 7 and 14. Values were normalized for the control mean for each experiment. Data presented as mean±SEM. One-way ANOVA, uncorrected Fisher’s LSD (*p<0.05). N=3-4 independent cultures. Abbreviations: CTRL – control; BDNF – Brain-derived neurotrophic factor

## Notes

### Competing Interest Statement

The authors have declared no competing interest.

